# Mitochondrial stress-induced GDF15-GFRAL axis promotes anxiety-like behavior and CRH-dependent anorexia

**DOI:** 10.1101/2021.09.22.461199

**Authors:** Carla Igual Gil, Bethany M. Coull, Wenke Jonas, Rachel Lippert, Mario Ost, Susanne Klaus

**Affiliations:** Department of Physiology of Energy Metabolism, German Institute of Human Nutrition Potsdam-Rehbruecke (DIfE), 14558 Nuthetal, Germany; Institute of Nutritional Science, University of Potsdam, 14469 Potsdam, Germany; Department of Neurocircuit Development and Function, German Institute of Human Nutrition; Department of Experimental Diabetology, German Institute of Human Nutrition Potsdam-Rehbruecke (DIfE), 14558 Nuthetal, Germany; German Center for Diabetes Research, 85764 München-Neuherberg, Germany; Institute of Anatomy, University of Leipzig, 04103, Leipzig, Germany; NeuroCure Cluster of Excellence, Charité Universitätsmedizin, Berlin, Germany

## Abstract

Growth differentiation factor 15 (GDF15) is a stress-induced cytokine that modulates food intake and energy metabolism. Until now, most mechanistic studies on GDF15 rely on pharmacological interventions using exogenous GDF15, but little is known about its mode of action when induced both chronically and endogenously. Mitochondrial stress is one of the most described physiological conditions that induces GDF15^1^, and therefore an important model to study the underlying mechanisms of endogenous GDF15’s action. Here, using a mouse model of mitochondrial dysfunction via elevated respiratory uncoupling in skeletal muscle, we show a circadian oscillation of muscle-derived GDF15 to promote a daytime-restricted anorexia without signs of nausea or reduced physical activity, contrary to findings using recombinant GDF15^2–5^. We find that mitochondrial stress-induced GDF15 associates with increased anxiety and hypothalamic corticotropin releasing hormone (CRH) induction, without further activation of the hypothalamic-pituitary-adrenal (HPA) axis and corticosterone response. Strikingly, the daytime-restricted anorexia, lean phenotype, systemic shift in substrate metabolism and anxiety-like behavior are completely abolished in conditions of mitochondrial stress coupled with genetic ablation of the GDF15 receptor GDNF receptor alpha-like (GFRAL), which is predominantly expressed in the hindbrain. Finally, we demonstrate that stress-induced GDF15-GFRAL signaling is required for hypothalamic CRH induction to control diurnal food intake in a CRH-receptor 1 (CRHR1)-dependent manner. With this, we uncover for the first time a molecular target of the GDF15-GFRAL axis that links anxiolytic and anorectic behavior as downstream effects of the chronic activation of this pathway by mitochondrial stress.

## Main

GDF15 is acknowledged as a stress-induced cytokine which can be expressed and secreted by multiple tissues^6^. While the anorexigenic action of exogenous GDF15 is well described^7–10^, less is known about the endocrine targets and mechanisms governing endogenous stress-induced GDF15. The unique receptor for GDF15 is GDNF receptor alpha-like (GFRAL) which is only expressed in the hindbrain (area postrema, AP, and nucleus of the solitary tract, NTS) and signals through the tyrosine kinase co-receptor Ret^7–10^. Increasing evidence suggests that the central GDF15-GFRAL axis acts as a mediator of the physiological responses to visceral malaise states such as the induction of food aversion, nausea and emesis which are preceding the onset of anorexia as shown in different animal models^2–4^. This fits with the well-known actions of the hindbrain AP in the control of nausea and vomiting^11^, beside its role as a control center of food intake^12^. Furthermore, recent data indicate that administration of recombinant GDF15 substantially lowers voluntary wheel running activity in mice^5^. Importantly, all evidence to date for the induction of malaise states through the GDF15-GFRAL pathway utilized supraphysiological doses of exogenously administered GDF15 or acute activation of the GDF15-GFRAL axis. Along these lines, little is known about the mode of action and physiological consequences of endogenously elevated GDF15, which would represent the conditions seen in physiologically relevant settings of disease. In tumor-bearing mice, antibody-mediated inhibition of GDF15-GFRAL activity reverses cancer cachexia^13^. In mice with chronically increased mitochondrial proteostatic stress, endocrine signaling of GDF15 was shown to regulate systemic energy expenditure^14–16^. Moreover, we have recently demonstrated the importance of mitochondrial-stress induced GDF15 to promote a diurnal, daytime restricted anorectic response to control systemic energy metabolism^17^. However, in order to understand the underlying mechanisms of its anorectic response, most studies used recombinant GDF15 or acute toxin treatments leading to short-term supraphysiological GDF15 levels while the mode of action of chronic, endogenous activation of the GFRAL-GDF15 pathway, representing a more pathophysiological state, has not been addressed so far. Using a mitochondrial dysfunction mouse model (HSA-*Ucp1*-TG [TG] mice)^18^ with chronically elevated circulating GDF15 levels, we here conducted a comprehensive in vivo behavioral and metabolic phenotyping in a sex- and GFRAL-dependent manner.

First, we established the diurnal profile of skeletal muscle *Gdf15* gene expression and circulating levels of wildtype (WT) and TG mice (**Fig. 1a-d**). In both male and female TG mice, *Gdf15* mRNA expression showed a diurnal oscillation (**Fig. 1a,c**) which was mirrored by the plasma GDF15 concentrations (**Fig. 1b,d**) with a delay of approximately 4h between gene expression and secretion. In order to determine whether nausea or pica behavior could be a consequence of endogenous elevated GDF15, we provided mice with a choice of chow and kaolin (a nonfood substance which rodents consume in response to emetogenic stimuli) and recorded their respective consumption over two consecutive days. Maximum daytime GDF15 levels in TG mice coincided with an almost complete avoidance of food intake, resulting in significantly lower daytime food intake compared to WT (**Fig. 1e,f**) in line with our previous findings^17^. Importantly, this daytime anorexia via mitochondrial stress-induced GDF15 was not linked to an induction of nausea or pica behavior. Both male and female TG mice showed no differences in the amount and pattern of kaolin consumption which was negligible during the daytime (**Fig. 1g,h**). Furthermore, in line with our previous data on normal voluntary wheel running activity of TG mice^17,19^ and contrary to recent findings on pharmacological GDF15 treatment^5^, we here show that physical cage activity reflected the regular nocturnal murine activity pattern, and daytime as well as total activity was similar in WT and TG (**Fig. 1i,j**), supporting the lack of malaise in TG mice.

**Figure 1.**
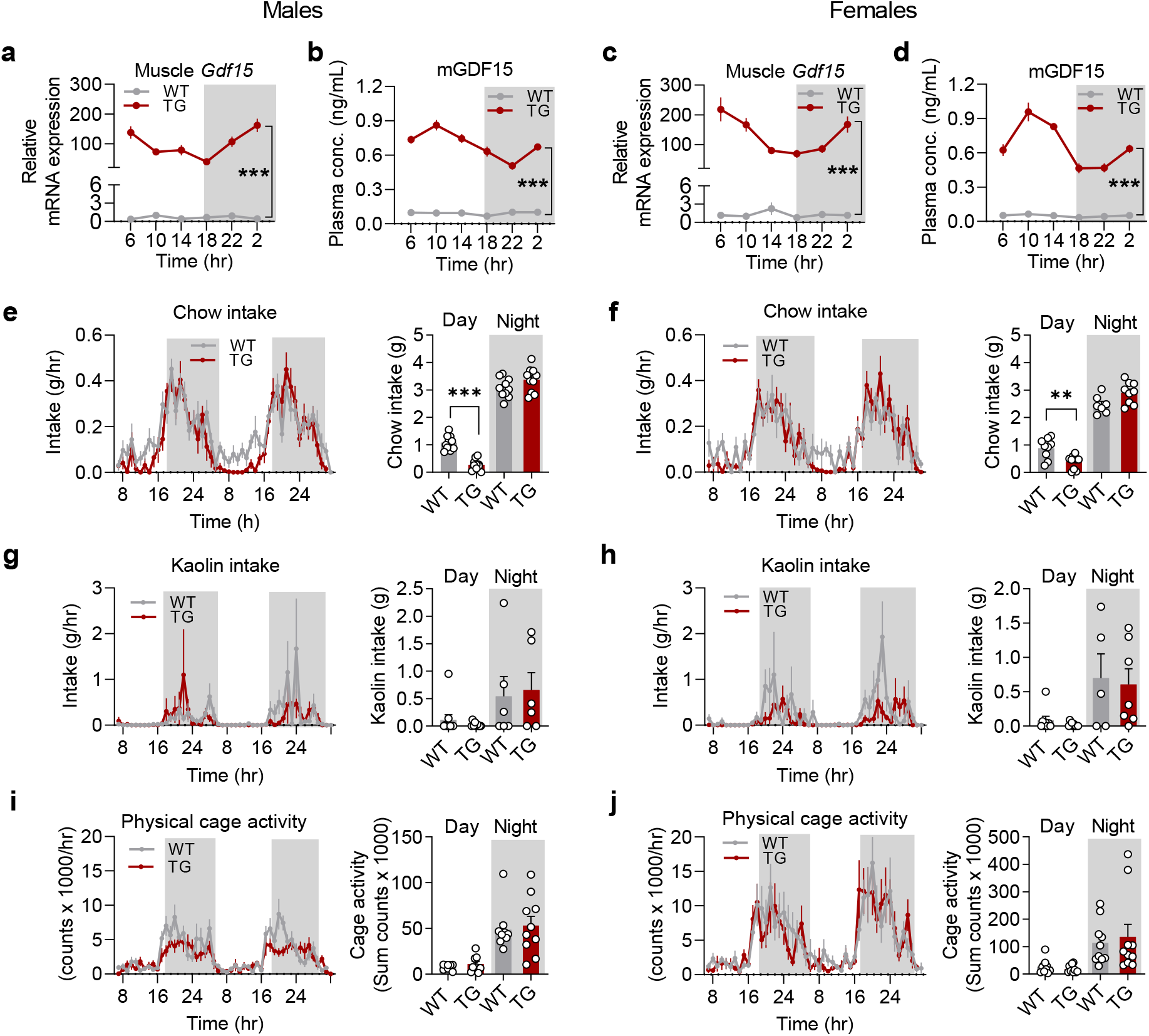
Mitochondrial stress-induced daytime anorexia via GDF15 is not associated with nausea or reduced physical activity. **a, c**, Circadian pattern of *Gdf15* mRNA expression in quadriceps muscle (n=7-8). **b, d,** circulating GDF15 plasma levels (n=9-10). **e, f,** Daily pattern and total chow (n=10). **g, h,** kaolin intake during a kaolin pica preference test (n=10). **i, j,** Daily pattern and total physical cage activity counts. The right and left panel of the figure correspond to male and female mice, respectively. Data correspond to wildtype (WT) and HSA-*Ucp1*-TG (TG) mice. Data are presented as mean +/− SEM,**P<0.01; ***P<0.001

Interestingly, genetic ablation of GDF15 in mice showed a decreased anxiety and increased exploratory behavior^20^, suggesting that physiological levels of GDF15 might be effective in increasing anxiety and stress related behavior. For exploration of anxiogenic effects upon mitochondrial stress, we further conducted open field (OFT) and elevated plus maze (EPM) tests to asses anxiety and exploratory behavior in TG mice. These tests were performed between 9 and 11hr, i.e. at the time of the GDF15 plasma peak in TG mice (**Fig. 1b,d**). In the OFT, TG mice spent more time in the corners and showed less time and entries into the center (**Fig. 2a,b**) suggestive of a decreased exploratory behavior. TG mice also showed an increased anxiety-like behavior compared to WT mice as evidenced by an increased freezing time during the EPM (**Fig. 2c,e**). Of note, male but not female TG mice showed a higher induction of circulating corticosterone after the EPM than wildtype littermates, suggestive of an increased HPA-axis responsiveness (**Fig. 2d,f**). These data indicate that mitochondrial stress and chronically elevated GDF15 associate with anxiety-like behavior and decrease exploratory behavior in mice.

**Figure 2.**
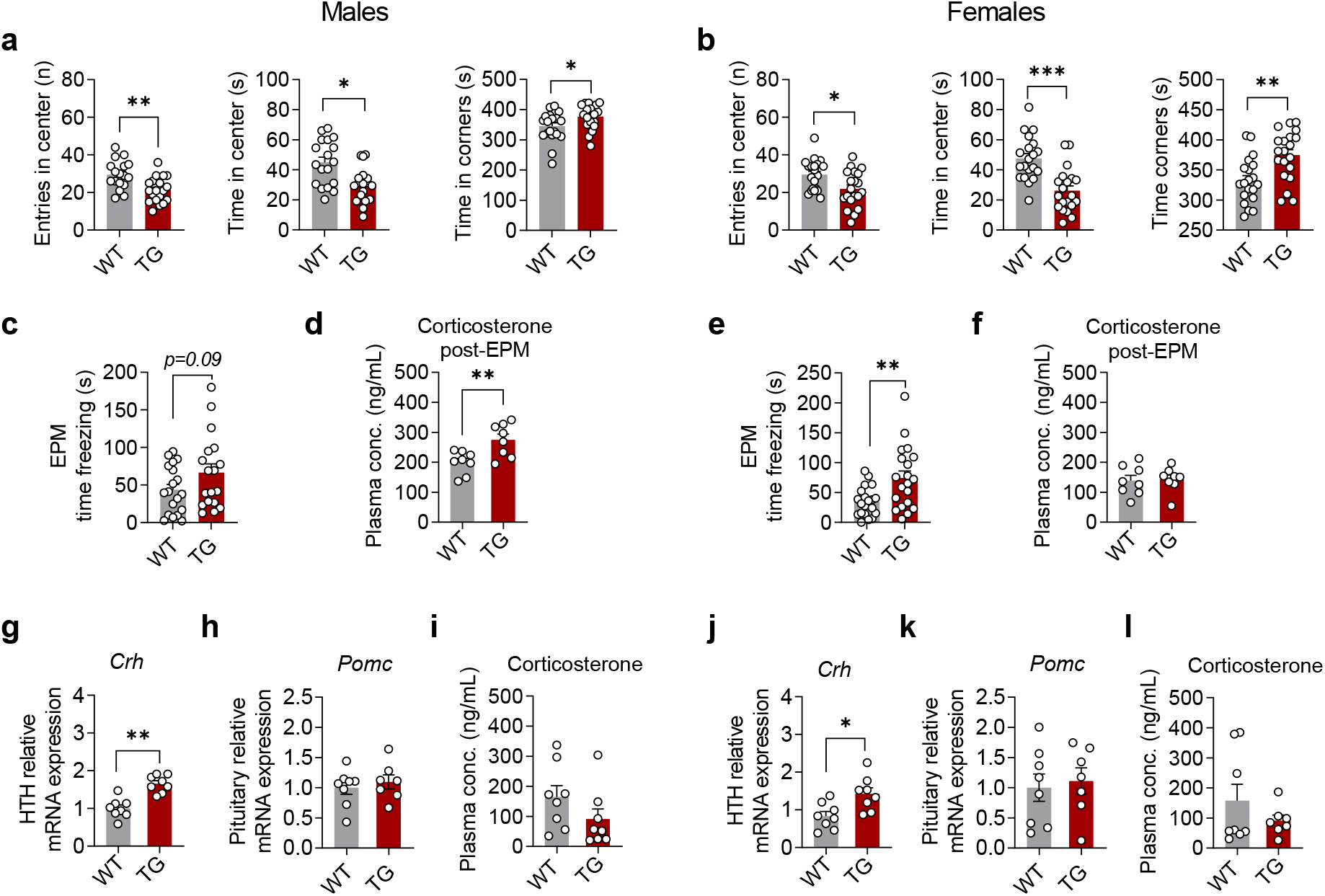
Mitochondrial stress associates with increased anxiety-like behavior and hypothalamic Crh induction. **a, b,** Entries in the center, time spent in the center and time spent in the corners during an open field test (n=20). **c, e** Time spent freezing during an elevated plus maze (EPM) test (n=20). **d, f,** post-EPM plasma corticosterone levels (n=8). **g, j,** Hypothalamus (HTH) *Crh* relative mRNA expression (n=8). **h, k,** Pituitary *Pomc* relative mRNA expression (n=8). **i, l,** Basal plasma corticosterone concentration of mice sacrificed at 10hr (n=7-8). The right and left panel of the figure correspond to male and female mice, respectively. Data correspond to wildtype (WT) and HSA-*Ucp1*-TG (TG) mice. Data are presented as mean +/− SEM, *P<0.05; **P<0.01; ***P<0.001

Activation of the GDF15-GFRAL pathway by recombinant GDF15 leads to increased cFOS in the lateral parabrachial nucleus (PBN), the oval sub-nucleus of the bed nucleus of the stria terminalis (ovBNST), the central amygdala (CeA) and the paraventricular nucleus of the hypothalamus (PVH)^21^. Interestingly, the PVH is the site of release of corticotropin-releasing hormone (CRH), the first step in the activation of the HPA-axis, a crucial regulator of the stress response known to be mediated via hindbrain neuron activation. CRH release from PVH neurons triggers the systemic release of adrenocorticotropin (ACTH) from the pituitary gland resulting in an increased adrenal secretion of glucocorticoids^22^. Therefore, we hypothesized that the GDF15-GFRAL axis might modulate stress related anxiety as well as ingestive behavior through the activation of the HPA-axis. First, to assess a possible activation of the HPA stress axis in TG mice we analyzed hypothalamic *Crh* expression in the PVH from mice dissected at 10hr. Indeed, *Crh* gene expression was significantly increased in both male and female TG mice compared to WT mice (**Fig. 2g,j**). Interestingly, pituitary expression of *Pomc*, the gene encoding ACTH, was also not increased in TG mice (**Fig. 2h,k**). Moreover, plasma corticosterone levels in undisturbed mice were not increased in TG mice at the time of highest circulating GDF15 (10hr) (**Fig. 2i,l**).

Next, in order to evaluate the importance of the GDF15-GFRAL pathway for the modulation of ingestive- and stress-related behavior we generated GFRAL-knockout male and female TG mice (TGxGfKO), which were investigated at 17-20 weeks of age in comparison with WT and GFRAL knockout (GfKO) mice. GFRAL ablation, as evidenced by non-detectable *Gfral* mRNA in the hindbrain (**Fig. 3a,d**), did not affect muscle *Gdf15* expression but led to slightly increased circulating GDF15 in male TG mice (**Fig. 3b,c**). In female TG mice muscle *Gdf15* expression was slightly increased by loss of GFRAL while circulating GDF15 was not affected (**Fig. 3e,f**).

**Figure 3.**
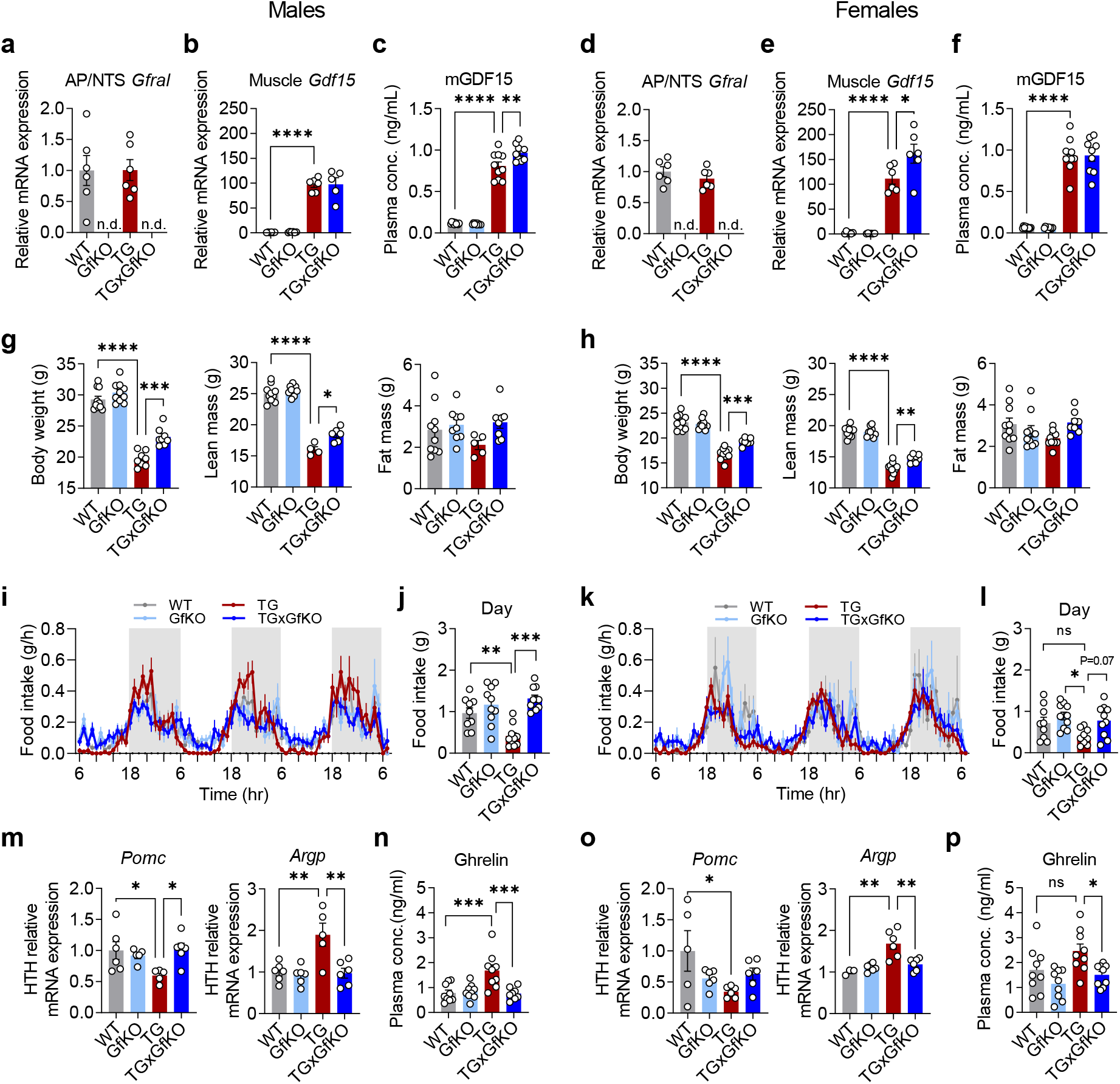
Stress-induced GDF15-GFRAL axis is responsible for lean phenotype and marked daytime anorexia. **a, d,** Gfral mRNA expression in the area postrema (AP) and nucleus of the solitary tract (NTS) (n=6). **b, e,** Muscle *Gdf15* mRNA expression (n=6). **c, f,** Circulating GDF15 plasma levels (n=10). **g, h,** Body weight and body composition (lean and fat mass) (n=10). **i, j, k, l,** Longitudinal assessment of food intake during three consecutive days (n=10). **k, m,** Quantification of daytime food intake (n=10). **m, o,** Hypothalamus (HTH) Pomc and Agrp gene expression (n=6). **n, p,** Plasma level of total ghrelin at daytime (10am) (n=9). The right and left panel of the figure correspond to male and female mice, respectively. Data correspond to wildtype (WT), *Gfral*-KO (GfKO), HSA-*Ucp1*-TG (TG) and HSA-Ucp1-TGx*Gfral*-KO (TGxGfKO) mice. Data are presented as mean +/− SEM, *P<0.05; **P<0.01; ***P<0.001; ****P<0.0001

In line with our previous data^23^, the body weight of TG mice was reduced due to a substantially lower lean mass. This was partially restored by GFRAL ablation which significantly increased body weight and lean mass of male and female TGxGfKO in comparison to TG mice (**Fig. 3g, h**). The muscle atrophy of TG mice, as evidenced by largely reduced quadriceps weight was not affected by GFRAL ablation, while the weight of other organs and tissues such as liver, heart, and different fat depots was increased to in TGxGfKO compared to TG mice (**Fig. S1a-d**). This suggests that muscle wasting of TG mice is a direct consequence of their muscle mitochondrial dysfunction and independent of the adaptive modulation of body composition which is a consequence of the GDF15-GFRAL pathway activation. Of note, GFRAL ablation alone had no effect on any of the assessed phenotypic traits in GfKO mice.

Next, we evaluated the relevance of GDF15-GFRAL axis activation for the shift in ingestive behavior induced by endogenously elevated GDF15 using high resolution metabolic phenotyping (**Fig. 3i,k**). Both in male and female mice, daytime anorexia was completely prevented by GFRAL ablation (**Fig. 3i,j,k,l**), although this effect was more pronounced in male mice. Interestingly, male TG mice showed a nighttime increase in food intake that was also GFRAL dependent (**Fig. 3i, Fig S2 a**), while nighttime food intake in female mice was not affected in any group (**Fig. 3k, Fig S2 c**). The daily distribution of water intake reflected the food intake pattern (**Fig. S2e,g**), and there were no differences in cage activity between the groups (**Fig. S2f,h**). In order to examine whether the daytime-restricted anorexia was orchestrated by the known central appetite drivers, we measured hypothalamic *Pomc* and *Agrp* gene expression. While *Pomc* expression was reduced, *Agrp* was increased in TG mice which was normalized by GFRAL ablation (**Fig. 3m,o**). Moreover, plasma concentrations of total ghrelin, a well-described orexigenic hormone^24^, were increased in male and female TG mice but blunted in TGxGfKO mice (**Fig. 3n,p**). Overall, the pattern in TG mice reflects a state of negative energy balance and increased appetite^25^. Strikingly, TG mice show a paradoxically reduced food intake that, together with the classical markers mentioned above, resembles anorexia like phenotypes and suggests that the GDF15-GFRAL axis works through an alternative pathway that overrides the classic hypothalamic food intake regulation system. A consequence of the diurnal variation of energy balance in TG mice is an increased systemic metabolic flexibility as evident by an increased amplitude of the respiratory quotient (RQ)^17^. Ablation of GFRAL completely abolished the increased metabolic flexibility of male and female TG mice, resulting in a RQ pattern and amplitude similar to WT and GfKO mice (**Fig. S2b,d**).

Finally, we assessed the involvement of the GDF15-GFRAL pathway in the modulation of anxiety-like behavior and hypothalamic CRH induction. The increased anxiety-like phenotype of male and tendency in female TG mice (decreased entries into the center during OFT and increased freezing time in EPM) was abolished in TGxGfKO mice (**Fig. 4a,b,d,e**). The higher post-EPM plasma corticosterone of male TG mice compared to WT was absent after GFRAL ablation (**Fig. 4c**), while female mice again showed no differences in post-EPM corticosterone (**Fig. 4f**). Importantly, the increased hypothalamic *Crh* gene expression of TG mice was normalized by GFRAL ablation in both male and female mice (**Fig. 4 g,j**), indicating that GFRAL is completely responsible for its upregulation. In line with the results from the mouse study presented in Fig. 2, pituitary *Pomc* expression (**Fig. 4 h,k**) and circulating corticosterone (**Fig. 4i,l**) were similar in all genotypes confirming that there was no downstream activation of the HPA axis in TG mice. Interestingly, CRH receptor 1 (CRHR1) knockout mice show a nearly opposite phenotype compared to TG mice in relation to ingestive and anxiety-like behavior. They display an impaired stress response and reduced anxiety^26,27^ together with a light-phase-restricted increase in food intake^28^. This suggests that elevated GDF15-GFRAL signaling may work, in part, via downstream CRH-CRHR1 pathways to elicit the marked changes in behavior shown here. To test this, we used antalarmin, a specific antagonist of CRHR1^29^, to assess the importance of CRH signaling for the GDF15-GFRAL dependent diurnal anorexia. Intraperitoneal injection of antalarmin abolished the day time anorexia of male TG mice (**Fig. 4m**) demonstrating the CRH-CRHR1 dependency of the diurnal anorexia induced by chronic activation of the GDF15-GFRAL pathway (**Fig. 4n**). Despite the strong hypothalamic *Crh* induction (**Fig. 4j**), antalarmin injection did not affect food intake in female TG mice, highlighting the complexity of sex-differences in regulation of food intake, which requires further investigations in the future.

**Figure 4.**
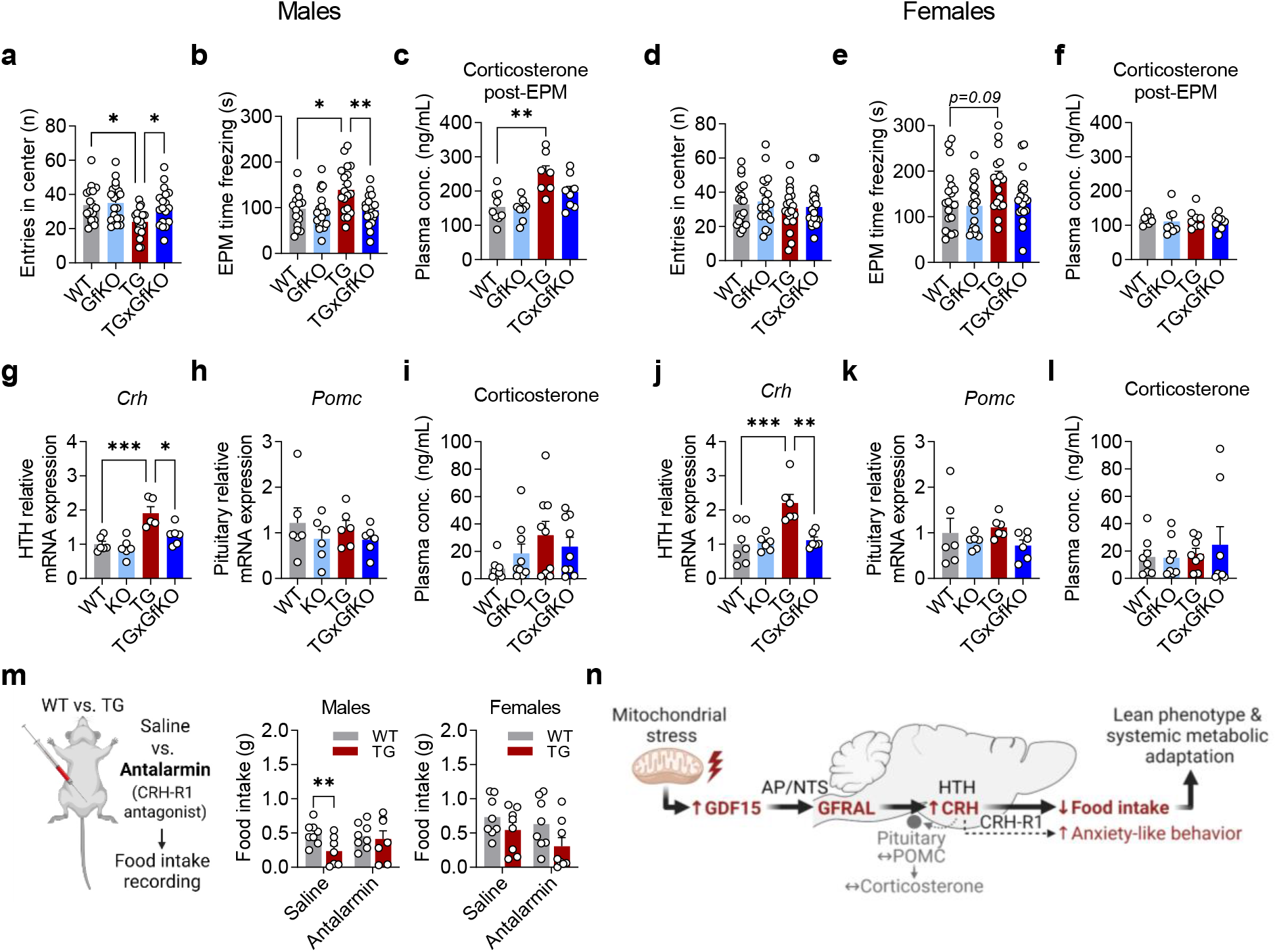
Stress-induced GDF15-GFRAL axis is required for the induction of anxiety-like behavior and hypothalamic CRH signaling to control daytime anorexia. **a, d,** Entries in the center during an open field test (n=20). **b, e,** Time spent freezing during an elevated plus maze (EPM) test (n=17-20). **c, f,** Plasma corticosterone levels after the EPM (n=8-9). **g, j,** Hypothalamus (HTH) *Crh* mRNA expression (n=8-9). **h, k**, Pituitary *Pomc* mRNA expression (n=6). **i, l,** Plasma corticosterone levels under routine conditions (n=8-9).. The right and left panel of the figure correspond to male and female mice, respectively. **m,** Daytime food intake after intraperitoneal injection of antalarmin, a selective CRHR1 antagonist, in male and female WT versus TG mice (n=8). **n,** Summary graph depicting the suggested central mechanism leading to modulation of ingestive and anxiety behavior by the GDF15-GFRAL pathway. Data correspond to wildtype (WT), *Gfral*-KO (GfKO), HSA-*Ucp1*-TG (TG) and HSA-*Ucp1*-TGx*Gfral*-KO (TGxGfKO) mice. Data are presented as mean +/− SEM, *P<0.05; **P<0.01; ***P<0.001. Cartoons in **m+n** were created with BioRender.

The importance of the GDF15-GFRAL pathway in appetite regulation has emerged only recently owing to the identification of GFRAL as the unique receptor for GDF15 and its localization exclusively in the hindbrain^30^. While investigations so far focused on anorexia induced by acute activation of the GDF15-GFRAL pathway, we reveal here that physiological, chronic activation of this pathway by endogenously elevated GDF15 induces hypothalamic CRH, increases anxiety-like behavior, and modulates the diurnal regulation of ingestive behavior in a sex-specific manner.

It was already shown in 2007 that GDF15 injection leads to activation of hindbrain neurons as well as in hypothalamic areas involved in appetite regulation such as the PVH^31^, but the nature of the GFRAL neurons in the hindbrain and their neural connections are just starting to be elucidated. Several studies connected GFRAL-expressing AP/NTS neurons to a neuronal circuitry which induces nausea and associated behavioral responses to different visceral malaise-inducing stimuli^3,21,32^. Thus, there seems to be a general consensus that the anorexia induced by acute activation of the GDF15-GFRAL pathway represents a sickness-behavior. Our data based on chronic activation of the GDF15-GFRAL pathway by lifelong elevated GDF15 levels challenges this notion. While we indeed find that GDF15-GFRAL signaling leads to a daytime-restricted anorexia via increased hypothalamic CRH mediated by CRHR1 activation, we find no evidence that this is sickness-related. In this line, it was recently shown that the physiological induction of endogenous circulating GDF15 by a prolonged strenuous exercise did not induce exercise-related food aversion or acute suppression of food intake in contrast to pharmacological application of GDF15^5^. Interestingly, single-nucleus RNA sequencing revealed that GFRAL-expressing neurons co-express the glucagon-like peptide 1 receptor (GLP1R)^32^. The most common side effect of GLP1R agonists used as anti-diabetic drugs in humans is the induction of nausea^33^. Therefore, it could be speculated that the over-activation of GFRAL/GLP1R neurons by supraphysiological doses of GDF15 leads to nausea as a secondary side effect.

Our data show clearly that the regulation of ingestive behavior by endogenous activation of the GDF15-GFRAL pathway is rather complex, therefore opening a number of avenues of research. *Gdf15* is a clock controlled gene^34,35^ and diurnal oscillations in circulation levels have been reported in healthy humans^5,36^. Considering the reversed circadian activity pattern in humans and mice, the diurnal pattern GDF15 in TG mice reflects the human pattern. The implications of this circadian/diurnal rhythmicity for the role of GDF15 in appetite control, especially in patho-physiological states have not yet been addressed. To date, most studies on the GDF15-GFRAL axis regulation of food intake have shown a clear, immediate anorectic response after activation of this pathway. Nevertheless, none of these studies considered the circadian patterns of the predominantly nocturnal food intake in rodents. Rather, they assessed ingestive behavior at single timepoints during the light-phase. Here we show that, while the activation of the GDF15-GFRAL pathway leads to a daytime-restricted anorexia in both male and female mice, it also increases food intake at nighttime in male mice, indicating that the GDF15-GFRAL axis not only has an anorectic role, but it rather induces a shift in ingestive behavior comprising both anorectic and orexigenic pathways. In goldfish, GDF15 was found to dose dependently increase food intake, thus acting as an orexigenic rather than an anorectic mediator^37^ supporting the notion that the GDF15-GFRAL pathway plays a complex role in ingestive behavior beyond the induction of anorexia and sickness behavior.

We further demonstrate that chronic activation of the GDF15-GFRAL axis in TG mice leads to an induction of hypothalamic *Crh*, without activation of the HPA axis. Only in male TG mice the stress induced increase in corticosterone was augmented. It was recently shown that an acute elevation of endogenous or recombinant GDF15 in mice and rats leads to an activation of the HPA axis evidenced by highly increased corticosterone levels 4 to 6 hours after treatment^38^. While our data clearly support the notion of an induction of hypothalamic CRH by GDF15-GFRAL signaling, our mouse model of chronic elevated GDF15 shows that chronic activation of the GDF15-GFRAL pathway might increase stress susceptibility but does not induce a sustained increase of corticosterone levels. Importantly, while acute administration of recombinant GDF15 promotes supraphysiological plasma GDF15 levels of >30ng/ml after 1-hr and >10ng/ml 4-hrs post -treatment in mice^38^, circulating GDF15 levels in TG mice oscillate in a range of 0.35 – 1.4ng/ml (Fig. 1b,d) in a circadian manner. This might explain differences in HPA axis induction and highlights the importance of studying animal models with chronic, endogenous elevated GDF15 in order to understand its biological action beyond canonical pharmacological effects.

Hypothalamic CRH has long been recognized as a catabolic mediator, suppressing food intake in animals and humans^39,40^. CRH effects on appetite and satiety are largely mediated by CRHR1 activation^41^. For the first time, we show here that chronic activation of the GDF15-GFRAL pathway acts via CRHR1 signalling to drive a diurnal anorexia, which is independent of the induction of visceral malaise or the HPA axis. There is increasing evidence that CRH neurons of the PVH are central players not only in appetite regulation but also in linking stress and anxiety behavior^42^. Here we uncover CRH as a key downstream player of the GDF15-GFRAL axis in the modulation of food intake and anxiety-like behavior. Nevertheless, further studies will be required to determine how GDF15-GFRAL-induced hypothalamic CRH signaling through CRHR1 modulates both anxiolytic and anorectic behavior and which brain regions are involved in this complex physiological response.

## Methods

### Animals

Mice with a C57BL/6J background were used for all experiments. *Gfral* heterozygous mice were purchased from Mutant Mouse Regional Resource Centers (MMRRC) and back-crossed to a C57BL/6J background. To generate our experimental genotypes, HSA-*Ucp1*-transgenic mice (TG) were crossed with *Gfral*-heterozygous (Gf-het) mice to obtain TGxGf-het mice, which were then further crossed with Gf-het mice obtaining wild-type (WT), *Gfral*-KO (GfKO), TG, and TGx*Gfral*-KO (TGxGfKO) mice. Mice were fed a standard chow diet (Sniff, Soest, Germany) with *ad libitum* access. All mice were kept group-housed and random-caged until sacrifice at 20 weeks of age, when organs were collected. For circadian studies of GDF15 oscillations mice were single-caged one week prior to sacrifice at 17 weeks of age. Unless otherwise indicated, animals were sacrificed between 9 and 11hr. All animal experiments were approved by the ethics committee of the Ministry of Agriculture and Environment (State Brandenburg, Germany, permission number 2347-16-2020).

### Behavioral testing

The Open Field Test (OFT) and the Elevated Plus Maze (EPM) test were performed in the same animals aged around 10 and 12 weeks, respectively, for a duration of 10min between 9 and 11hr. The open field apparatus consisted of a 50×50cm enclosure. The mouse was placed in the center of the field and recorded with a camera using the software ANY-maze 5.2. The Elevated Plus Maze apparatus consisted of two open (30×5×0.5cm) and two closed (30×5×15cm) arms, which cross each other in a middle platform (5×5×0.5cm) designated as the center. To start the test mice were placed in one of the open arms and were recorded using ANY-maze 5.2.

### *In vivo* metabolic phenotyping and kaolin preference test

Body composition was measured with quantitative magnetic resonance (QMR, EchoMRI 2012 Body Composition Analyzer, Houston, USA). The respiratory quotient (RQ=CO_2_ produced/O_2_ consumed) was measured by indirect calorimetry with simultaneous recording of cage activity, food and water intake (TSE PhenoMaster, TSE Systems, Germany). This system was also used for the assessment of kaolin intake during a kaolin preference test. While in the indirect calorimetry cages, mice were presented with two food hoppers, one with standard chow diet and the other with kaolin pellets (Research Diets, #K50001). Food intake of both food hoppers was recorded during 48 hours, with a change of the position of the food hoppers after the first 24 hours.

### Gene expression analysis

RNA was isolated with a phenol-chlorophorm-based extraction using peqGOLD Trifast (VWR, #732-3314) followed by a DNase digest (Fischer Scientific, #EN0521). Synthesis of cDNA was performed with the LunaScript RT SuperMix Kit (NEB, #E3010L). For quantitative real-time PCR (qPCR) analyses, 5ng of cDNA, LUNA Universal Probe qPCR Mastermix (NEB, #M3004E) and 1,5 μM of primers in a total volume of 5μl were used. Measurements were performed on a ViiA™ 7 Real-Time PCR System from Applied Biosystems. The following primer sequences were used: *Gdf15*: 5’ GAGCTACGGGGTCGCTTC 3’ (F), 5’ GGGACCCCAATCTCACCT 3’ (R); *Crh*: 5’ CAACCTCAGCCGGTTCTGAT 3’ (F), 5’ CAGCGGGACTTCTGTTGAGA 3’ (R); *Ucp1*: 5’ TGGAGGTGTGGCAGTATTC 3’ (F), 5’ AGCTCTGTACAGTTGATGATGAC 3’ (R); *Gfral*: 5’ CGAAATGATGAATTATGCAGGA 3’ (F), 5’ TGCAGGTCTCATCTTCATGG 3’ (R); *Pomc*: 5’ AACCTGCTGGCTTGCATC 3’ (F), 5’ GACCCATGACGTACTTCCG 3’ (R); *Agrp*: 5’ TTGGCGGAGGTGCTAGAT 3’ (F), 5’ ACTCGTGCAGCCTTACACAG 3’ (R).

### Plasma analyses

Whole blood was collected through heart puncture in heparin tubes (Sarstedt, #41.1503.005), centrifuged at 9,000 g for 10min at 4°C and plasma was stored at −80°C. For measurement of routine and post-EPM corticosterone levels blood was collected from the tail vein. Plasma GDF15 was quantified using the Mouse/Rat GDF-15 Quantikine ELISA Kit (Bio-techne, #MGD150). Plasma corticosterone was measured with a Corticosterone ELISA kit (Enzo, #ADI-900-097). Plasma ghrelin was measured with Meso-Scale Discovery (MSD) multiplex assay (MSD instruments, USA).

### Antalarmin treatment and food intake recording

To perform the antalarmin treatment experiment with simultaneous recording of food intake a Food Access System (Phenomaster, TSE Systems GmbH) was used. This system allows for automatic recording of food intake. Animals were single-caged in the Food Access System cages for 72 hours prior to start of the measurements. The measurements took place during the next three consecutive days. On day one, no intervention was performed in order to obtain basal food intake measurements. On day two, an intraperitoneal (IP) injection of vehicle (0,9% saline and 10% Cremophore® EL (Merk, #238470)) was conducted. On day three, the mice received an intraperitoneal injection of the CRHR1 antagonist antalarmin (30mg/kg) (Sigma-Aldrich, #A8727). IP injections were conducted at 8hr. Food intake was recorded until 24 hours after vehicle and antalarmin injection, respectively.

### Statistical analysis

Statistical analyses were performed using GraphPad Prism 9 (GraphPad Software, San Diego, CA, USA). All data are expressed as mean with SEM. Data were tested for normality using D’Agostino & Pearson normality test. According to the group number, a Student’s t-test (unpaired, 2-tailed) or a one-way ANOVA followed by the Tukey’s multiple comparison test was used to determine differences between genotypes. At *P <* 0.05 statistical difference was assumed and denoted by \**P <* 0.05, \*\**P <* 0.01, \*\*\**P <* 0.001, \*\*\*\**P <* 0.0001. Statistically significant differences are only shown for WT vs. TG and TG vs. TGxGfKO for comprehensive reasons. In addition to individual data, data are shown as mean +/− SEM.

## Data availability

All data that support the findings of this study are available from the corresponding author upon reasonable request.

## Acknowledgements

This work was supported by grants from the German Research Foundation (DFG, grant no. Kl 613/23– 1) and NutriAct (Research Stimulus Grant to CIG). The authors would like to thank Carolin Borchert, Petra Albrecht and Antje Sylvester for excellent technical assistance. The cartoons in Figure 4 were created with BioRender.

---

**Figure S1.**
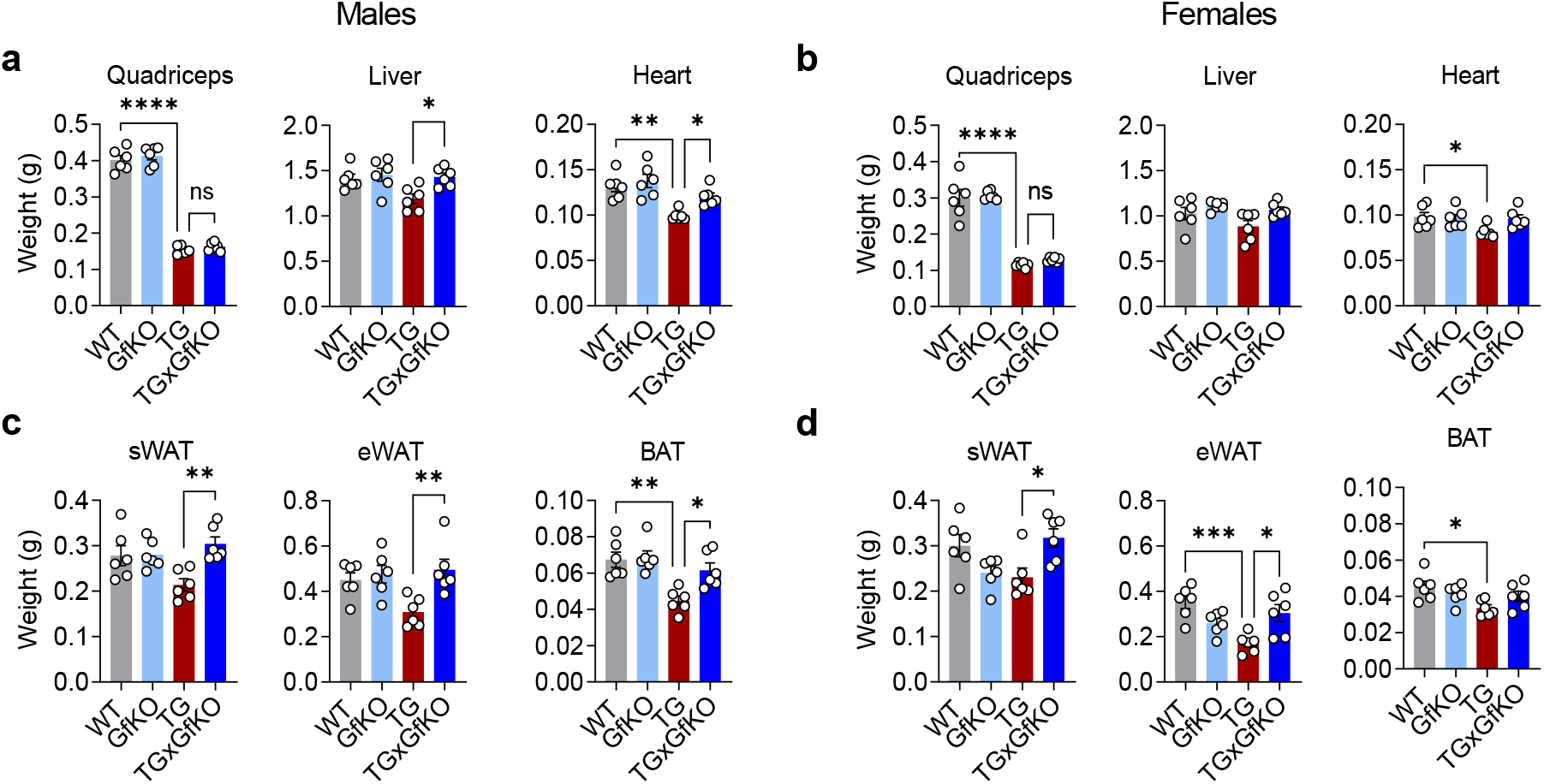
Loss of GFRAL increases lean and fat tissue weight upon mitochondrial stress in TG mice. **a, b,** Quadriceps, liver and heart tissue weight (n=6). **c, d,** Subcutaneous adipose tissue (sWAT), epididymal adipose tissue (eWAT) and brown adipose tissue (BAT) weights (n=6). The right and left panel of the figure correspond to male and female mice, respectively. Data correspond to wildtype (WT), *Gfral*-KO (GfKO), HSA-*Ucp1*-TG (TG) and HSA-*Ucp1*-TGx*Gfral*-KO (TGxGfKO) mice. Data are presented as mean +/− SEM, *P<0.05; **P<0.01; ***P<0.001; ****P<0.0001.

**Figure S2.**
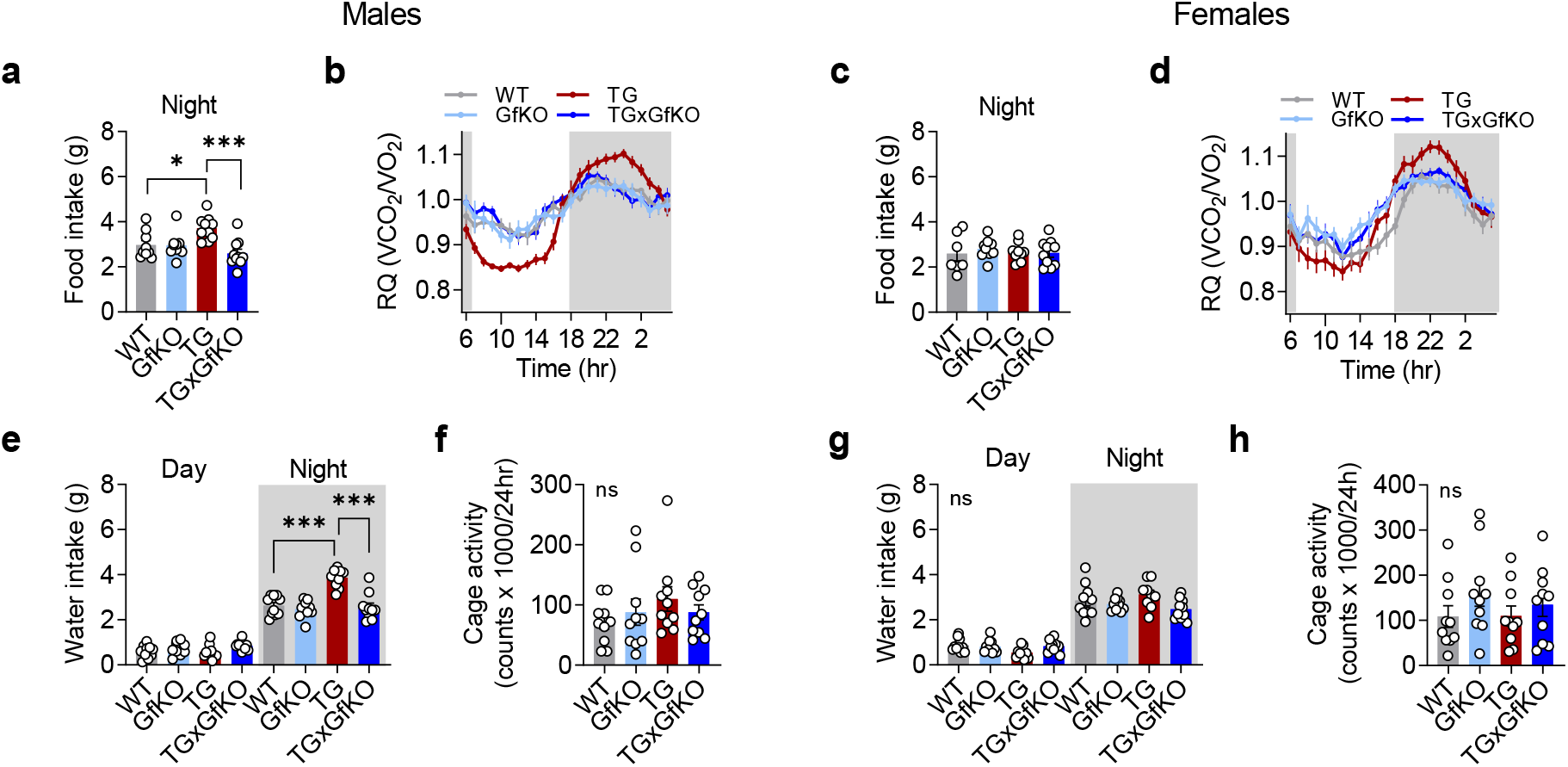
GFRAL is involved in a nighttime shift in ingestive behavior in male but not in female mice. **a, c,** Quantification of total night time food intake (n=10). **b, d,** Respiratory quotient (RQ) shown hourly over 24 hours (n=10). **e, g,** Day and night water intake (n=10). **f, h,** Physical cage activity shown hourly over 24 hours (n=10). The right and left panel of the figure correspond to male and female mice, respectively. Data correspond to wildtype (WT), *Gfral*-KO (GfKO), HSA-*Ucp1*-TG (TG) and HSA-*Ucp1*-TGx*Gfral*-KO (TGxGfKO) mice. Data are presented as mean +/− SEM,*P<0.05; ***P<0.001.

## References

1. Keipert, S. & Ost, M. Stress-induced FGF21 and GDF15 in obesity and obesity resistance. Trends in Endocrinology & Metabolism, doi:https://doi.org/10.1016/j.tem.2021.08.008 (2021).

2. Borner, T. et al. GDF15 Induces Anorexia through Nausea and Emesis. Cell Metab 31, 351–362 e355, doi:10.1016/j.cmet.2019.12.004 (2020).

3. Sabatini, P. V. et al. GFRAL-expressing neurons suppress food intake via aversive pathways. Proc Natl Acad Sci U S A 118, doi:10.1073/pnas.2021357118 (2021).

4. Borner, T. et al. GDF15 Induces an Aversive Visceral Malaise State that Drives Anorexia and Weight Loss. Cell Rep 31, 107543, doi:10.1016/j.celrep.2020.107543 (2020).

5. Klein, A. B. et al. Pharmacological but not physiological GDF15 suppresses feeding and the motivation to exercise. Nat Commun 12, 1041, doi:10.1038/s41467-021-21309-x (2021).

6. Lockhart, S. M., Saudek, V. & O’Rahilly, S. GDF15: A Hormone Conveying Somatic Distress to the Brain. Endocrine Reviews 41, 610–642, doi:10.1210/endrev/bnaa007 (2020).

7. Hsu, J. Y. et al. Non-homeostatic body weight regulation through a brainstem-restricted receptor for GDF15. Nature 550, 255–259, doi:10.1038/nature24042 (2017).

8. Emmerson, P. J. et al. The metabolic effects of GDF15 are mediated by the orphan receptor GFRAL. Nat Med 23, 1215–1219, doi:10.1038/nm.4393 (2017).

9. Mullican, S. E. et al. GFRAL is the receptor for GDF15 and the ligand promotes weight loss in mice and nonhuman primates. Nat Med 23, 1150–1157, doi:10.1038/nm.4392 (2017).

10. Yang, L. et al. GFRAL is the receptor for GDF15 and is required for the anti-obesity effects of the ligand. Nat Med 23, 1158–1166, doi:10.1038/nm.4394 (2017).

11. Miller, A. D. & Leslie, R. A. The area postrema and vomiting. Front Neuroendocrinol 15, 301–320, doi:10.1006/frne.1994.1012 (1994).

12. Hyde, T. M. & Miselis, R. R. Effects of area postrema/caudal medial nucleus of solitary tract lesions on food intake and body weight. Am J Physiol 244, R577–587, doi:10.1152/ajpregu.1983.244.4.R577 (1983).

13. Suriben, R. et al. Antibody-mediated inhibition of GDF15-GFRAL activity reverses cancer cachexia in mice. Nat Med, doi:10.1038/s41591-020-0945-x (2020).

14. Choi, M. J. et al. An adipocyte-specific defect in oxidative phosphorylation increases systemic energy expenditure and protects against diet-induced obesity in mouse models. Diabetologia 63, 837–852, doi:10.1007/s00125-019-05082-7 (2020).

15. Chung, H. K. et al. Growth differentiation factor 15 is a myomitokine governing systemic energy homeostasis. J Cell Biol 216, 149–165, doi:10.1083/jcb.201607110 (2017).

16. Kang, S. G. et al. Differential roles of GDF15 and FGF21 in systemic metabolic adaptation to the mitochondrial integrated stress response. iScience 24, 102181, doi:10.1016/j.isci.2021.102181 (2021).

17. Ost, M. et al. Muscle-derived GDF15 drives diurnal anorexia and systemic metabolic remodeling during mitochondrial stress. EMBO Rep, e48804, doi:10.15252/embr.201948804 (2020).

18. Klaus, S., Rudolph, B., Dohrmann, C. & Wehr, R. Expression of uncoupling protein 1 in skeletal muscle decreases muscle energy efficiency and affects thermoregulation and substrate oxidation. Physiol Genomics 21, 193–200, doi:10.1152/physiolgenomics.00299.2004 (2005).

19. Coleman, V. et al. Partial involvement of Nrf2 in skeletal muscle mitohormesis as an adaptive response to mitochondrial uncoupling. Sci Rep 8, 2446 (2018).

20. Low, J. K. et al. First Behavioural Characterisation of a Knockout Mouse Model for the Transforming Growth Factor (TGF)-beta Superfamily Cytokine, MIC-1/GDF15. PLoS One 12, e0168416, doi:10.1371/journal.pone.0168416 (2017).

21. Worth, A. A. et al. The cytokine GDF15 signals through a population of brainstem cholecystokinin neurons to mediate anorectic signalling. Elife 9, doi:10.7554/eLife.55164 (2020).

22. Herman, J. P. et al. Regulation of the Hypothalamic-Pituitary-Adrenocortical Stress Response. Compr Physiol 6, 603–621, doi:10.1002/cphy.c150015 (2016).

23. Keipert, S., Voigt, A. & Klaus, S. Dietary effects on body composition, glucose metabolism, and longevity are modulated by skeletal muscle mitochondrial uncoupling in mice. Aging cell 10, 122–136, doi:10.1111/j.1474-9726.2010.00648.x (2011).

24. Inui, A. Ghrelin: an orexigenic and somatotrophic signal from the stomach. Nat Rev Neurosci 2, 551–560, doi:10.1038/35086018 (2001).

25. Quarta, C. et al. POMC neuronal heterogeneity in energy balance and beyond: an integrated view. Nat Metab 3, 299–308, doi:10.1038/s42255-021-00345-3 (2021).

26. Smith, G. W. et al. Corticotropin releasing factor receptor 1-deficient mice display decreased anxiety, impaired stress response, and aberrant neuroendocrine development. Neuron 20, 1093–1102, doi:10.1016/s0896-6273(00)80491-2 (1998).

27. Timpl, P. et al. Impaired stress response and reduced anxiety in mice lacking a functional corticotropin-releasing hormone receptor 1. Nat Genet 19, 162–166, doi:10.1038/520 (1998).

28. Muller, M. B., Keck, M. E., Zimmermann, S., Holsboer, F. & Wurst, W. Disruption of feeding behavior in CRH receptor 1-deficient mice is dependent on glucocorticoids. Neuroreport 11, 1963–1966, doi:10.1097/00001756-200006260-00031 (2000).

29. Lowery-Gionta, E. G. et al. Corticotropin releasing factor signaling in the central amygdala is recruited during binge-like ethanol consumption in C57BL/6J mice. J Neurosci 32, 3405–3413, doi:10.1523/JNEUROSCI.6256-11.2012 (2012).

30. Breit, S. N., Brown, D. A. & Tsai, V. W. The GDF15-GFRAL Pathway in Health and Metabolic Disease: Friend or Foe? Annu Rev Physiol 83, 127–151, doi:10.1146/annurev-physiol-022020-045449 (2021).

31. Johnen, H. et al. Tumor-induced anorexia and weight loss are mediated by the TGF-beta superfamily cytokine MIC-1. Nat Med 13, 1333–1340, doi:10.1038/nm1677 (2007).

32. Zhang, C. et al. Area Postrema Cell Types that Mediate Nausea-Associated Behaviors. Neuron, doi:10.1016/j.neuron.2020.11.010 (2020).

33. Filippatos, T. D., Panagiotopoulou, T. V. & Elisaf, M. S. Adverse Effects of GLP-1 Receptor Agonists. Rev Diabet Stud 11, 202–230, doi:10.1900/RDS.2014.11.202 (2014).

34. Tasaki, H. et al. Profiling of circadian genes expressed in the uterus endometrial stromal cells of pregnant rats as revealed by DNA microarray coupled with RNA interference. Front Endocrinol (Lausanne) 4, 82, doi:10.3389/fendo.2013.00082 (2013).

35. Zhao, L. et al. The nuclear receptor REV-ERBalpha represses the transcription of growth/differentiation factor 10 and 15 genes in rat endometrium stromal cells. Physiol Rep 4, doi:10.14814/phy2.12663 (2016).

36. Tsai, V. W. et al. Serum Levels of Human MIC-1/GDF15 Vary in a Diurnal Pattern, Do Not Display a Profile Suggestive of a Satiety Factor and Are Related to BMI. PLoS One 10, e0133362, doi:10.1371/journal.pone.0133362 (2015).

37. Blanco, A. M., Bertucci, J. I., Velasco, C. & Unniappan, S. Growth differentiation factor 15 (GDF-15) is a novel orexigen in fish. Mol Cell Endocrinol 505, 110720, doi:10.1016/j.mce.2020.110720 (2020).

38. Cimino, I. et al. Activation of the hypothalamic-pituitary-adrenal axis by exogenous and endogenous GDF15. Proc Natl Acad Sci U S A 118, doi:10.1073/pnas.2106868118 (2021).

39. Rothwell, N. J. Central effects of CRF on metabolism and energy balance. Neurosci Biobehav Rev 14, 263–271, doi:10.1016/s0149-7634(05)80037-5 (1990).

40. Richard, D. & Baraboi, D. Circuitries involved in the control of energy homeostasis and the hypothalamic-pituitary-adrenal axis activity. Treat Endocrinol 3, 269–277, doi:10.2165/00024677-200403050-00001 (2004).

41. Lemos, J. C. et al. Severe stress switches CRF action in the nucleus accumbens from appetitive to aversive. Nature 490, 402–406, doi:10.1038/nature11436 (2012).

42. Daviu, N., Bruchas, M. R., Moghaddam, B., Sandi, C. & Beyeler, A. Neurobiological links between stress and anxiety. Neurobiol Stress 11, 100191, doi:10.1016/j.ynstr.2019.100191 (2019).

